# The primary afferent activity cannot capture the dynamical features of muscle activity during reaching movements

**DOI:** 10.1101/138859

**Authors:** Russell L. Hardesty, Matthew T. Boots, Sergiy Yakovenko, Valeriya Gritsenko

## Abstract

The stabilizing role of sensory feedback in relation to realistic 3-dimensional movement dynamics remains poorly understood. The objective of this study was to quantify how primary afferent activity contributes to shaping muscle activity patterns during reaching movements. To achieve this objective, we designed a virtual reality task that guided healthy human subjects through a set of planar reaching movements with controlled kinematic and dynamic conditions that minimized inter-subject variability. Next, we integrated human upper-limb models of musculoskeletal dynamics and proprioception to analyze motion and major muscle activation patterns during these tasks. We recorded electromyographic and motion-capture data and used the integrated model to simulate joint kinematics, joint torques due to muscle contractions, muscle length changes, and simulated primary afferent feedback. The parameters of the primary afferent model were altered systematically to evaluate the effect of fusimotor drive. The experimental and simulated data were analyzed with hierarchical clustering. We found that the muscle activity patterns contained flexible task-dependent groups that consisted of co-activating agonistic and antagonistic muscles that changed with the dynamics of the task. The activity of muscles spanning only the shoulder generally grouped into a proximal cluster, while the muscles spanning the wrist grouped into a distal cluster. The bifunctional muscle spanning the shoulder and elbow were flexibly grouped with either proximal or distal cluster based on the dynamical requirements of the task. The composition and activation of these groups reflected the relative contribution of active and passive forces to each motion. In contrast, the simulated primary afferent feedback was most related to joint kinematics rather than dynamics, even though the primary afferent models had nonlinear dynamical components and variable fusimotor drive. Simulated physiological changes to the fusimotor drive were not sufficient to reproduce the dynamical features in muscle activity pattern. Altogether, these results suggest that sensory feedback signals are in a different domain from that of muscle activation signals. This indicates that to solve the neuromechanical problem, the central nervous system controls limb dynamics through task-dependent co-activation of muscles and non-linear modulation of monosynaptic primary afferent feedback.

**New & Noteworthy:** Here we answered the fundamental question in sensorimotor transformation of how primary afferent signals can contribute to the compensation for limb dynamics evident in muscle activity. We combined computational and experimental approaches to create a new experimental paradigm that challenges the nervous system with passive limb dynamics that either assists or resists the desired movement. We found that the active dynamical features present in muscle activity are unlikely to arise from direct feedback from primary afferents.

## Introduction

Movement is the product of interactions between neural control signals and the musculoskeletal dynamics that depends on limb anatomy (Yakovenko, 2011; Gritsenko et al., 2016). The motor control problem is then solved within a system with coupled neural and mechanical dynamical elements (Schoner and Kelso, 1988; Taga et al., 1991). Actions are generated not only by the internal signals within the central nervous system (CNS), but also by the compensatory actions arising from the intrinsic muscle properties that shape actuation through limb mechanics. These actions generate the task-dependent sensory feedback signals that in turn shape ongoing neural commands (Prochazka:2007ba; Ting et al., 2015). The nature of these neuromechanical interactions can be unraveled by examining the features contained within the kinematic and kinetic signals that describe the movement. For example, the kinetic signals are related to the active joint torques generated through muscle contractions to move the limb. These torques depend on the passive joint torques that arise from the interactions between internal and external forces, such as limb inertia, gravity, etc. The assistive or resistive contributions of these passive torques are present in all movements of segmented limbs. In a reaching motion, the simultaneous rotation of humerus and radius/ulna segments in the same direction in one plane causes resistive passive torques at shoulder and elbow joints that oppose the intended motion. These resistive passive torques are compensated for by active muscle torques at the appropriate joints (Almeida et al., 1995; Gribble and Ostry, 1999; Koshland et al., 2000; Pigeon et al., 2003; Debicki and Gribble, 2005; Kurtzer et al., 2008; Gritsenko et al., 2011). In contrast, assistive passive properties can be employed by the CNS to minimize effort (Debicki et al., 2011). The best example of the assistive action of gravity is in the locomotion of passive dynamic walkers, where the interaction torques generate knee flexion in early swing and gravity helps swing the foot forward (Collins et al., 2005). Skilled baseball players increase the efficiency of their throws by using the whiplash action of interaction torques at the elbow that are initiated by active torque applied at the shoulder (Hirashima et al., 2007). In contrast to movements with resistive passive torques, the activity of muscles is reduced to allow the assistive passive torques to assist with the movement. However, as with any positive feedback, the assistive action of passive torques may be destabilizing and difficult to control with low muscle contractions. This instability may be compensated for by the viscoelastic properties of muscles and spinal reflexes that can resist perturbations and ensure stable control with minimal supraspinal input (Asatryan and Feldman, 1965; Brown and Loeb, 2000; Yakovenko et al., 2004; Prochazka and Yakovenko, 2007; Valero-Cuevas et al., 2015). Other evidence suggests that coordinated co-contraction of antagonistic muscles driven by predictive descending signals can increase joint stiffness and reduce any instability (De Serres and Milner, 1991; Burdet et al., 2001; Gribble et al., 2003; Darainy et al., 2007).

The pattern of muscle modulation by the assistive and resistive limb dynamics during reaching remains sparsely described. This is due in part to the complexity of the sensorimotor control loop. The prevailing theory of motor control is that the CNS embeds complex mechanical limb dynamics in order to shape control signals and incorporate sensory feedback (Kluzik et al., 2008; Kurtzer et al., 2008; Wagner and Smith, 2008; Dideriksen et al., 2015). This embedding at the supra-spinal level is often referred to as internal models (Kawato et al., 1987), which can be observed, for example, as dynamical units in the primary motor cortex (Churchland et al., 2012). There are also strong arguments for the embedding of mechanical limb dynamics at the spinal level. For example, interaction torques can be counteracted through homonymous reflexes from muscle spindles and tendon organs of biarticular muscles that span the relevant joints (Lacquaniti et al., 1991; Prochazka et al., 1997; Verschueren et al., 1999). Additionally, interaction torques can be counteracted through heteronomous reflexes of monoarticular muscles that are coupled across relevant joints, as observed during locomotion in animals (Koshland and Smith, 1989; Abelew et al., 2000). However, the compensation for interaction torques caused by perturbations at the arm has been shown to occur at longer delays than expected from spinal reflexes (Kurtzer et al., 2009; 2014), and it requires the integration of proprioceptive inputs by the primary motor cortex (Pruszynski et al., 2011). Thus, it is still unclear how and how much proprioception contributes to the embedding of mechanical limb dynamics.

The contribution of sensory feedback to the ongoing motor activity can be modified through several neural pathways. Most relevant to the scope of the current study are the changes to the primary afferent activity caused by the fusimotor drive. The fusimotor drive is often one of the unknown variables; it is provided through the innervation of muscle spindles by dynamic and static γ motoneurons (Boyd and Gladden, 1985). During movement, the dynamic fusimotor action changes mainly the velocity sensitivity of the Ia primary afferents, while the static fusimotor action changes mainly their length sensitivity (Prochazka, 2011). How exactly the activity of γ motoneurons changes during reaching movements in humans is unknown. However, the effect the fusimotor drive has on shaping the muscle spindle output can be broadly classified based on whether the fusimotor drive is constant or changing during movement. The former is defined as “fusimotor set”, where γ motoneuron activity remains constant during a given movement, but its level changes between different types of movements in order to adjust the sensitivity of the muscle spindles to the anticipated demands of the task (Prochazka et al., 1985; Prochazka, 1986). Alternatively, the sensory feedback could be coupled to the ongoing motor activity via “α-γ coactivation”, where the sensitivity of muscle spindles is maintained during muscle shortening by coupling the activity of γ motoneurons to the activity of α motoneurons (Granit, 1970; Hagbarth, 1993). The distinctions between these alternative scenarios can be recreated using Ia primary afferent feedback models with parameters responsible for the sensitivity of feedback to muscle length and velocity changes (Prochazka and Gorassini, 1998; Mileusnic et al., 2006).

The rationale for this study was to quantify how muscle activity pattern reflects the kinetic requirements of a reaching task and to determine if the primary afferent feedback is congruent with this pattern or with the pattern of kinematic signals. We also determined if the fusimotor drive can increase the congruency between primary afferent signals and the kinetic task requirements.

## Materials and Methods

### Experimental design and human subjects

We recruited 9 healthy adults (5 males, 4 females; age, 24.3 ± 1.8 years; weight, 76.3 ± 14.5 kilograms) to perform reaching movements to visual targets in a virtual reality (VR) environment. All procedures were approved by the West Virginia University Institutional Review Board (Protocol #1309092800).

Subjects performed three reaching tasks in the VR environment (Oculus Rift; Fig. 1A). Pairs of visual targets defined the starting and goal target locations for each task, (Fig. 1B). To minimize inter-subject variability in angular kinematics, the locations of virtual targets were derived using planar trigonometry based on the lengths of subject’s arm and forearm segments and the specific combination of initial and final shoulder and elbow angles required to create 3 tasks described below. The pairs of starting and goal visual targets were shown in a random sequence. The cue to move was the change of target color from red to green. Each task was repeated 24 times.

**Figure 1.**
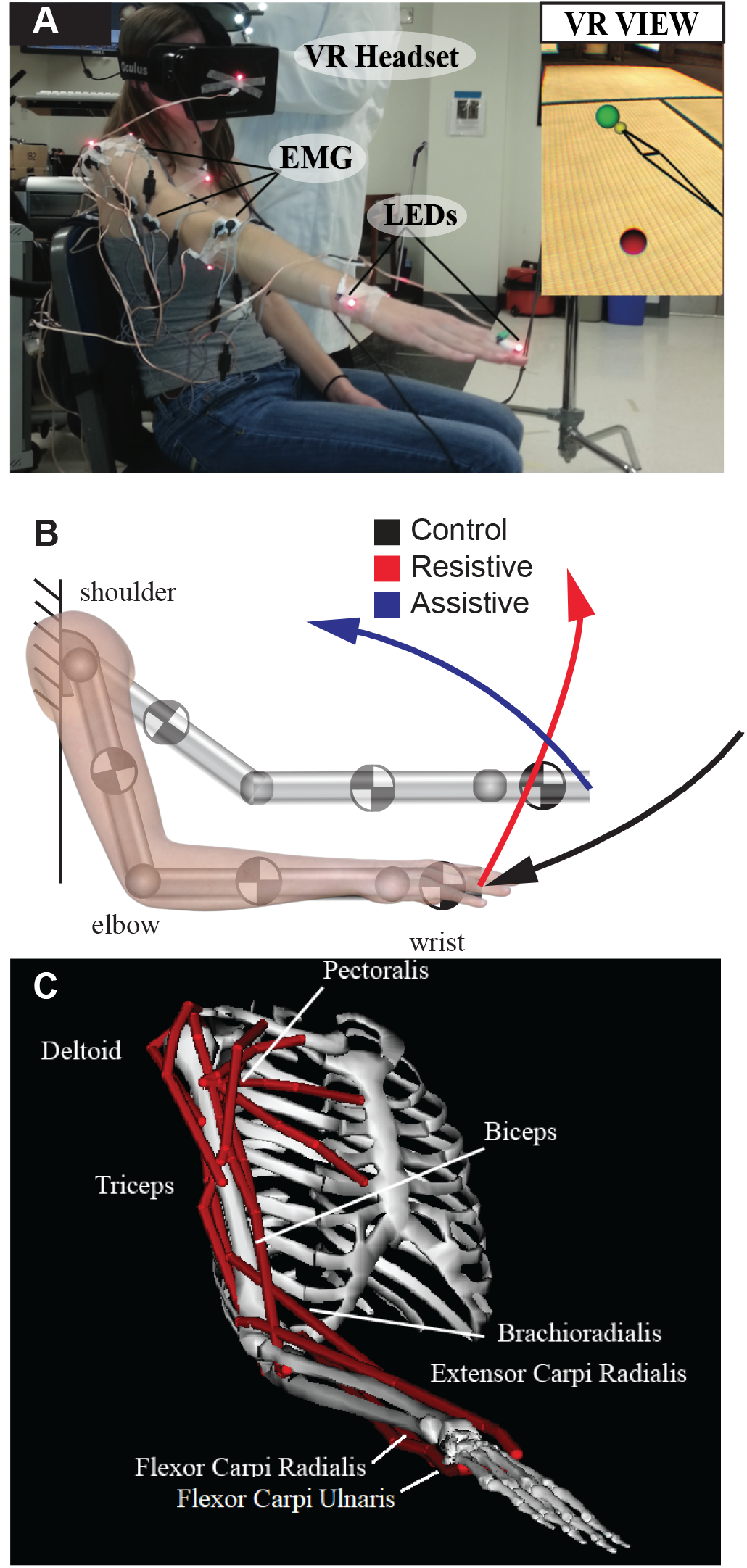
Illustrations of the experimental setup and arm models. A. Annotated photo of the setup; insert shows subject’s monocular view. Reaching target is in green, origin target is in red. Yellow sphere shows the location of subject’s fingertip and the black lines outline the major arm segments for visual feedback of arm location in VR. B. Colored lines show the fingertip trajectories of each of the three tasks. Arrows indicate the direction of motion toward the reaching target. The grey blocks show the locations and orientations of local coordinate systems used to obtain joint torques from motion capture. C. Illustration of the OpenSim model used to derive muscle lengths for the calculations of Ia primary afferent discharge. Red lines show the anatomical paths of each muscle from which EMG signals were recorded during experiments.

The virtual targets constrained endpoint trajectories to be in the same directions within the plane of movement across all subjects (Fig. 1B). During planar movement between these targets, the arm joints experienced different dynamical contexts, which defined tasks referred as Control, Resistive, and Assistive in the rest of the manuscript.

During the performance of each task, we recorded the kinematics of the shoulder, elbow, and wrist joints and electromyography (EMG) of 12 muscles that span those joints. The recorded muscles were the anterior and posterior deltoids (AD and PD, respectively), pectoralis major (Pec), teres major (TM), biceps brachii long and short heads (BicL and BicS, respectively), triceps brachii lateral and long heads (TriLa and TriLo, respectively), brachioradialis (Br), extensor carpi radialis (ECR), flexor carpi radialis (FCR), and flexor carpi ulnaris (FCU). These muscle abbreviations are used consistently throughout the manuscript and figures. Motion capture data were recorded at 480 hertz (Hz) using an Impulse system (PhaseSpace), and EMG signals were recorded at 2000 Hz with an MA400-28 system (MotionLab Systems). The start and end of each movement was defined by a jerk threshold method using wrist and elbow LED markers. The motion capture data were used to derive joint angles using linear algebra (Robertson et al., 2013). The EMG was processed consistent with SENIAM recommendations, it was high-pass filtered at 10 Hz, rectified, and low-pass filtered at 20 Hz. The resulting EMG profiles were averaged per task and normalized to the maximum across all tasks per subject.

### Ia model

To estimate the sensory contribution from muscle spindles during movement, we used Prochazka’s model of Ia primary afferent discharge (Prochazka, 1999), which offers a clear parametrization of static and dynamic responses. Although the model was derived from afferent recordings in cats, it was validated for humans by Malik et al. (2016). The spindle model relates afferent firing rate (*Ia*) to the time-varying muscle length (*l*) and its rate of change (*v*) as follows:

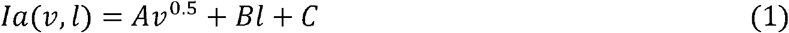

where the constant parameters (*A* = 65, *B* = 200, and *C* = 10) were validated empirically to reflect human microneurography data (Malik et al., 2016).

The changes of musculotendon length during movement were calculated in OpenSim using a modified musculoskeletal model of the human arm (Saul et al., 2015; Gritsenko et al., 2016) (Fig. 1C). This model was adjusted for each individual using subject’s morphology to scale bone segment and move proportionally origin and insertion for each muscle path. Muscle lengths were simulated by driving the adjusted model with the mean angular trajectories for each task and subject. The unit of parameter *A* and *B* in Eq.1 were normalized to a muscle rest length and its full physiological range of motion (ROM). The length was scaled so that extreme positions were 0 and 1. The velocity was normalized to the rest length per second units, where the rest length was defined as the length of the muscle in the middle of physiological ROM. The parameter space of A and B variations was explored in the context of variable fusimotor drive. The following parameter ranges were explored: *A*∈[33 200] and *B*∈[50 400], which resulted in 4 models referred to as follows: V33-L50, V33-L400, V200-L50, V200-L400, where V stands for velocity coefficient and L stands for muscle length coefficient.

Separately, we approximated α-γ coupling of both dynamic and static fusimotor inputs by incorporating recorded muscle activity profiles from the time-varying EMG (*a*) normalized to [0 1], which transformed equation (1) as follows:

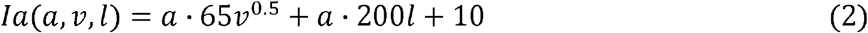

The model in (2) is referred to below as EMG-coupled Ia model.

For the regression analysis described below, the output of Eq.1-2 was normalized to the maximum of Ia discharge across all tasks per subject to obtain unitless values.

### Mechanical Coupling Between Muscles

In a previous study, we quantified the mechanical coupling between muscles based on their anatomy using a musculoskeletal model of the arm (Gritsenko et al., 2016). In that study, we used the same OpenSim model described above to obtain muscle lengths over the whole range of physiological joint postures. In the current study, we selected a subset of muscle length data from the Gritsenko et al. (2016) study for the muscles recorded here and included them in the hierarchal analysis described below. We used the muscle lengths obtained from the analysis across all postures rather than the muscle lengths calculated for the Ia modeling, because the former was calculated over a wider range of postures than the latter.

### Task dynamics

The dynamic mechanical model of a human upper-limb (Olesh et al., 2017) was used to compute joint torques from joint motions. The model was constructed in Simulink (MathWorks, Inc.); it comprised three segments and five DOFs, including the shoulder (flexion/extension, abduction/adduction, internal/external rotation), elbow (flexion/extension), and wrist (flexion/extension). The height and weight of each subject were used with anthropometric tables (Winter, 2009) to estimate the lengths and cylindrical inertias of the arm, forearm, and hand segments (Fig. 1B). To calculate active muscle torques, mean angular trajectories (Fig. 2A) from each subject and task were used to drive the subject-specific model in inverse dynamic simulations in Matlab (MathWorks, Inc.; Fig. 2B). The motion defined by our tasks was in the vertical plane. Subjects showed minimal out-of-plane motion as measured by angular trajectories about shoulder abduction/adduction and internal/external rotation DOFs. Therefore, only muscle torques about the shoulder flection/extension DOF was included in the analysis described below.

**Figure 2.**
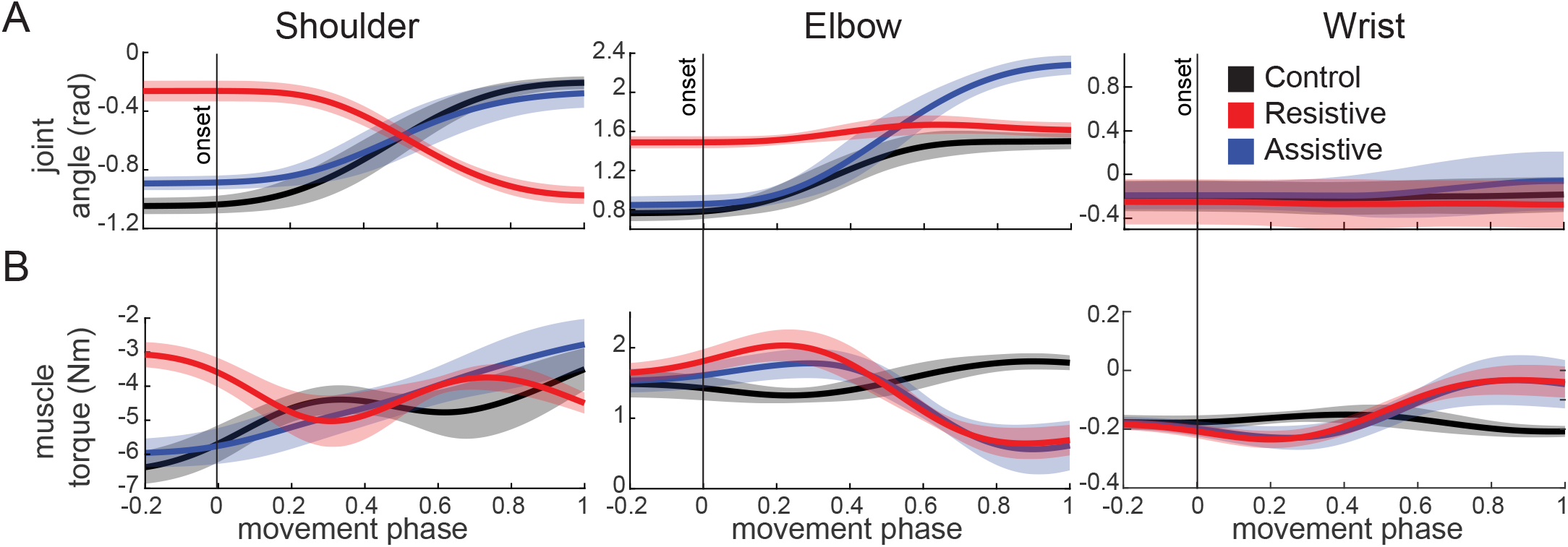
Signals calculated from motion capture. Thick lines show averages for each movement across all subjects, shaded areas show standard deviations across subjects. Only rotatum signals calculated for shoulder flexion/extension DOFs included in the following analyses are shown. Movement phase represents normalized duration of each movement with 0 indicating the start of movement (vertical onset line) and 1 indicating the end of movement. A. Joint angles; B. muscle torques; C. and D. flexion and extension rotatum signals respectively calculated from the muscle torques in B.

To describe the dynamics of each task, the three muscle torques about shoulder, elbow, and wrist were then used to calculate the following parameters. 1) The postural torque change was calculated as the difference between muscle torques averaged over 100 ms prior to onset and following the offset of movement. These postural torques are produced to maintain the arm in starting and final postures against the force of gravity. 2) The peak torque change in acceleration phase was calculated as the maximal change in torque between the start of movement to its halfway point. This measure is indicative of the amount of muscle force that is required to start the motion. 3) The mechanical muscle work was calculated by integrating a product between muscle torques and angular velocity as described in (Winter, 2009). When the direction of action matches between muscle torque and angular velocity, as indicated by the same sign (both positive or both negative), the mechanical muscle work is positive. This means that agonist muscle contractions about the corresponding DOF are concentric and they are actively producing the motion. When the direction of action is opposite between muscle torque and angular velocity, as indicated by opposite signs, the mechanical muscle work is negative. This means that agonist muscle contractions about the corresponding DOF are eccentric and the motion is produced by passive torques, such as gravity, interaction torques, etc. In our tasks, the wrist joint does not move, therefore the mechanical muscle work about the wrist is zero. This means that the muscle torque about the wrist reflects the isometric contraction of wrist and hand muscles that is required to stabilize the joint at a constant angle.

### Analysis

To quantify the common and distinct features in EMG and Ia profiles and to compare them to features obtained from muscle lengths we used hierarchical clustering of correlation matrix. All normalized profiles for each subject and each task were aligned on the onset of movement and normalized to its duration. The relationships between the EMG and Ia signals were characterized by a matrix of Pearson’s correlation coefficients (*r*). To reduce the probability of Type I errors, the *α* for determining the significance of *r* values was adjusted using the two-stage Benjamini, Krieger, and Yekutieli procedure for controlling false detection rate (Benjamini et al., 2006). The correlation matrix was then transformed into the heterogeneous variance explained (HVE) as follows:

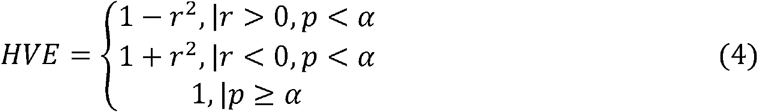

The HVE transforms the large positive *r* values that are characteristic of agonistic relationships into short distances close to 0 and the large negative *r* values corresponding to antagonistic relationships into long distances close to 2. To identify synergistic relationships between EMG and Ia, we applied hierarchical clustering to an unbiased HVE distance matrix using the linkage function with an unweighted average distance method (Gritsenko et al., 2016). The goodness of fit of the clustering model was assessed using the cophenetic correlation coefficient, which quantified how faithfully the hierarchical cluster tree represented the dissimilarities among observations. The magnitude of this value should be very close to 1 for a high-quality solution.

Clusters were compared using the Fowlkes-Mallows index (*B_k_*) to assess cluster similarity between separate hierarchical cluster trees (Fowlkes and Mallows, 1983). The Fowlkes-Mallows index represents a normalized number of common elements between clusters from different trees at the same cluster height. For example, *B_2_* indicates that the hierarchical trees were compared at the height, where only 2 clusters occur. Here, we explored *k* = [2, …, *n*], where *n* is half the number of signals being included in the hierarchical clustering. Thus, for two cluster trees with arbitrarily numbered clusters *i* = 1, …, *k* and *j* = 1, …, *k* we can use the number of objects between the *i*th cluster of one tree and *j*th cluster of the other tree (*m_ij_* to calculate the index as follows:

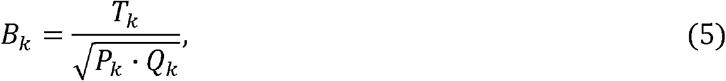

where

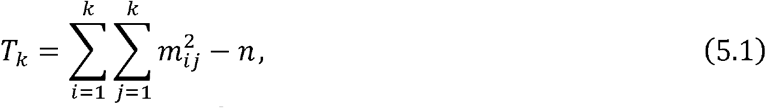

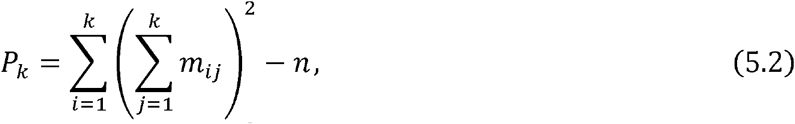

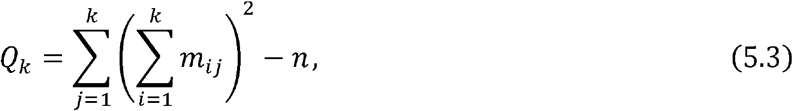

### Statistics

All values reported in results are means with standard deviations, unless stated otherwise. The shared variance between clusters defined by hierarchical clustering was assessed using t-tests. The t-tests were applied to squared correlation coefficients (R^2^) averaged across members of the cluster per subject per task. Subjects were assumed to represent independent samples. The combined p-values across subjects included in the tables were obtained using the Fisher’s combined probability test (Fisher, 1925).

The statistical comparison of hierarchical clustering between multiple signal modalities was based on permuting the hierarchical clustering trees to estimate the chance of observing spurious correlations. The hierarchal tree for each subject each movement type and each signal modality (Ia, EMG, muscle length) was randomly permuted 1000 times. Then the Fowlkes-Mallows index (B) was calculated between each of the permuted trees, which resulted in a population of *B* values that represents the distribution of noise. The distribution of experimental *B* values across tasks and subjects was compared to the corresponding noise distribution of *B* values to test the null hypothesis that *B_experimental_* = *B_noise_*. The p-value for each experimental B value was determined from the corresponding noise distribution for each subject using the percentile method (Efron and Tibshirani, 1993). This permutation analysis was applied to test two null hypotheses: 1) the similarity between Ia and EMG clusters is due to chance; 2) the similarity between Ia and muscle-length clusters is due to chance. Rejecting either of these null hypotheses means that the compared trees are similar to a greater extent than is expected by chance.

The statistical comparison of hierarchical clustering between tasks was based on bootstrapping the *B* values (Efron and Tibshirani, 1993). The *B* values for EMG clustering (*B_EMG_*) and Ia clustering (*B_Ia_*) calculated between tasks were resampled with replacement 1000 times. This resulted in two distributions of 45,000 *B_EMG_* and *B_Ia_* values for each task pair (Control-Resistive, Control-Assistive, and Resistive-Assistive). These data were used to test the null hypothesis that Ia clusters change between tasks the same way as the EMG clusters change between tasks. To test this hypothesis, we calculated the difference between the two distributions of *B_EMG_* and *B_Ia_* values, each comprising 1000 bootstraps per cluster number (*k = 2, …, 6*) per subject (N = 9). The p-value for each task pair was determined from the location of the 0 value in the resulting distribution of differences using the percentile method (Efron and Tibshirani, 1993).

The last set of null hypotheses addressed the extent to which the fusimotor drive can shape primary afferent discharge to capture the features of muscle activity pattern. The null hypothesis for each altered Ia model was that the similarity between Ia and EMG clusters is not increased by alternative fusimotor drives. To test this hypothesis the *B* values were calculated between the hierarchical clustering of Ia and EMG profiles for each of the models with altered coefficients (V33-L50, V33-L400, V200-L50, V200-L400, and EMG-coupled). To test the hypotheses, the distribution of *B* values from each of the altered model was subtracted from the corresponding *B* values based on Ia profiles from the Prochazka model. The p-value for each model with altered coefficients was determined from the location of the 0 value in the resulting distribution of differences using the percentile method (Efron and Tibshirani, 1993).

## Results

Subjects performed reaching tasks with consistent angular kinematics within the constraints defined by the VR targets. The angular excursions of each joint were similar across subjects for the three tasks (Fig. 2, top row). Because the subject’s movements were not restricted, most subjects moved slightly out of the sagittal plane and the experimental angular displacement differed from the predicted angles (Table 1). Despite slight out-of-plane motion, the VR targets defined the predicted tasks with distinct roles that limb dynamics played during movement, compensation for which was reflected in muscle torques. In the Control and Assistive tasks, the virtual targets defined joint excursions that required the shoulder and elbow joints to rotate in opposite directions. This caused assistive interaction torques between these joints similar to the Assistive task in (Gritsenko et al., 2011), which were associated with negative muscle work at the shoulder and positive muscle work at the elbow (Table 1). The sign of work indicates the direction of energy flow. The positive sign of work indicates concentric contractions that transfer energy from muscles to segments, while the negative sign of work indicates eccentric contractions during which the energy from external forces are overpowering the muscle action and doing the work (Winter, 2009). Thus, in the Control and Assistive tasks, the shoulder motion was largely passive, and the activity of shoulder muscles was compensating for external forces due to gravity and interaction torques. In the Control task, the elbow and wrist torques were the lowest across the three tasks (Fig. 2, black lines). The Assistive task was accompanied by decreasing postural torques in all joints, low acceleration shoulder torques, but high deceleration elbow and wrist torques (Fig. 2, blue lines; Table 1, third column). This shows that in the Assistive task most of the muscle action was to decelerate the limb accelerated primarily by the interaction torques and gravity. In contrast, in the Resistive the joint excursions were such that required the shoulder and elbow to rotate in the same direction, causing resistive interaction torques similar to the Resistive task in (Gritsenko et al., 2011). Altogether, this caused the opposite pattern of shoulder torques compared to that in the Control and Assistive tasks, while maintaining the same elbow and wrist torques to that in the Assistive task (Fig. 2 C and D). In the Resistive task, the mechanical muscle work was always positive, indicating that muscles contractions were concentric, and that the motion was produced with the least reliance on passive limb dynamics and gravity.

**Table 1.**
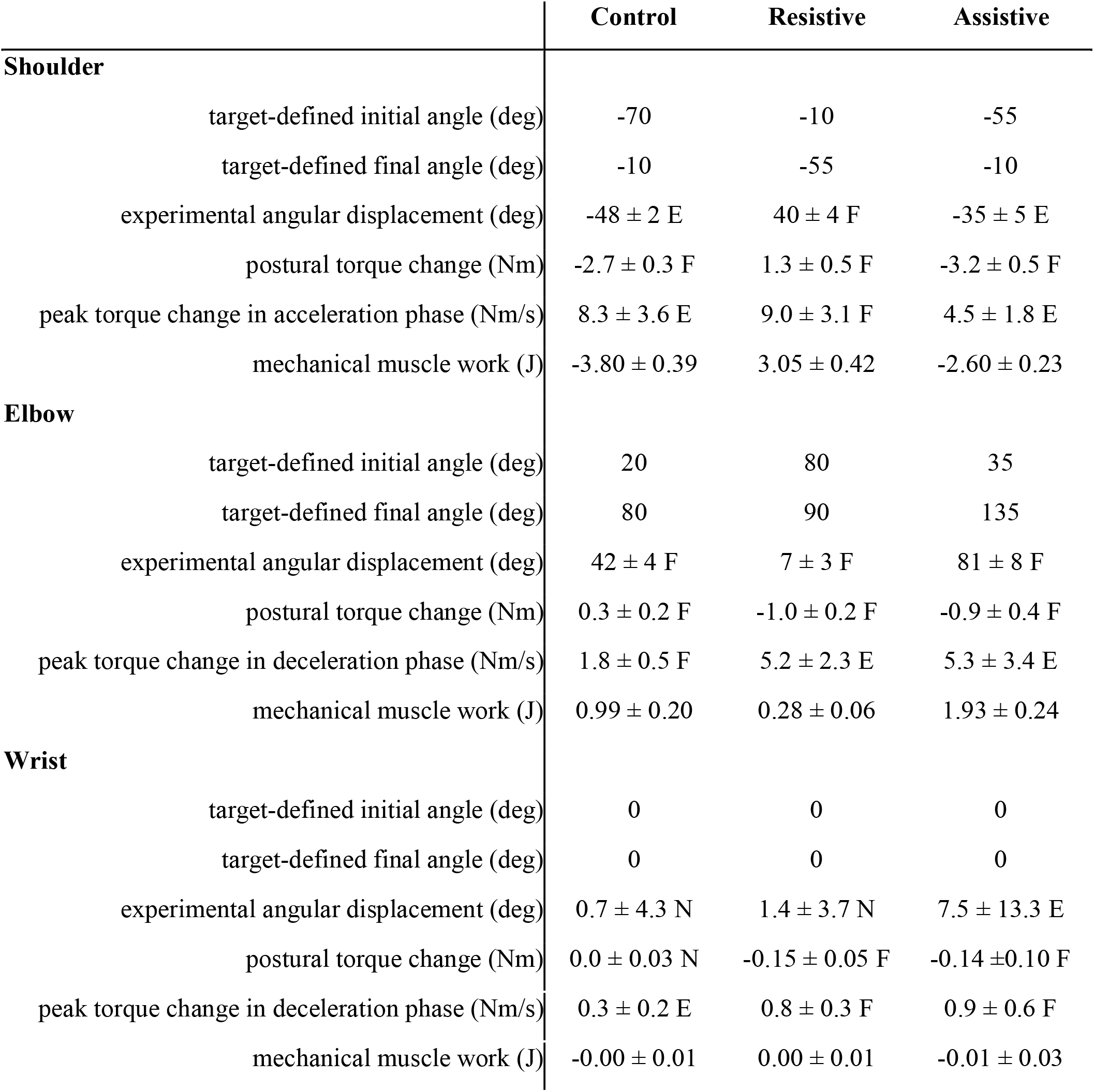
Table of task parameters. All joint angles are around flexion/extension axes of rotation for the three major joints included in the analysis. The sign of angles and torques indicates the direction of change of the corresponding measure, positive for increase and negative for decrease. Experimental values are averages with standard deviations across subjects. F indicates flexion direction of action; E indicates extension direction of action; N indicates no change. Negative values of mechanical muscle work imply work done by external forces, e.g. gravity and reaction from distal segments in the mechanical chain.

EMG and Ia profiles during movement reflected distinct aspects of the dynamical conditions set up by the tasks (Fig. 3). The opposite direction of shoulder motion in the Resistive task compared to the other two tasks was accompanied by alternating patterns of Ia signals in all four muscles that span only the shoulder joint, i.e. Pec, AD, PD, and TM. The EMG patterns of the corresponding muscles were less antagonistic, with the exception of the AD EMG profile (Fig. 3, second row). Furthermore, the EMG profiles of multiple muscles that span the elbow and wrist joints showed features that were not captured by the kinematics or dynamics of the tasks, such as increased activation of the TriLo, TriLa, BicL, BicS, Br, FCR, FCU, and ECR toward the end of movement in the Assistive task (Fig. 3, blue lines). This pattern was similar to the differences between tasks in the Ia profiles of the TriLo, TriLa, FCU, and FCR and opposite to that of the BicL, BicS, Br, and ECR.

**Figure 3.**
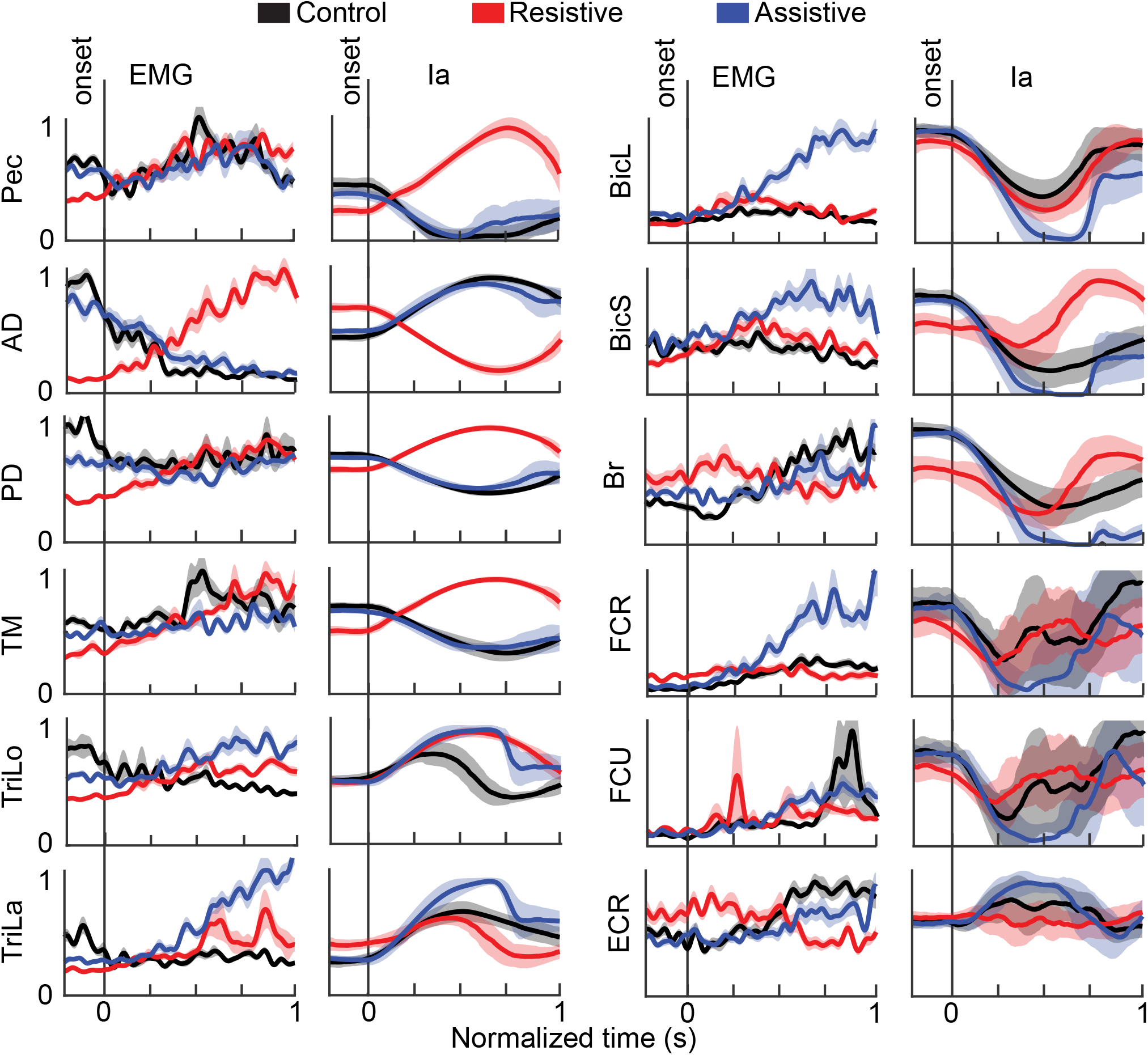
EMG and Ia profiles. Thick lines show averages for each movement across all subjects. Shaded areas show the standard error of the mean across subjects for EMG signals and standard deviation across subjects for Ia signals. Movement phase represents normalized duration of each movement as in Fig. 2.

Hierarchal cluster analysis has captured the relationships between EMG and Ia signals with high precision, as evidenced by high cophenetic coefficient of 0.81 ± 0.087. EMG profiles were found to contain different features that reflect the dynamical context of each task. In the Assistive task, muscle activity shared the most variance (mean R^2^ within the cluster = 0.36 ± 0.11 SD across subjects), indicating more co-contraction than in the other tasks (Control R^2^ = 0.20 ± 0.05, Resistive R^2^ = 0.26 ± 0.08). The differences in R^2^ values between the Assistive task and both the Control and Resistive task were significant (Assistive vs. Control t = 6.5, p < 0.001; Assistive vs. Resistive t = 2.96, p = 0.018). In the Assistive task, the co-contraction was evident in a single EMG cluster of TriLa/TriLo/BicL/BicS/Br/FCR/FCU/ECR that was present in 80 % of subjects (R^2^ = 0.48 ± 0.15; Fig. 4 Assistive). Proximal muscle activity clustered less frequently with lower shared variance than captured by the distal cluster (Pec/TM/AD/PD R^2^ = 0.20 ± 0.06). These data show that the largely passive shoulder motion and assistive passive forces in the Assistive task (Table 1) were accompanied by weak co-contraction of proximal muscles and high co-contraction of distal muscles.

**Figure 4.**
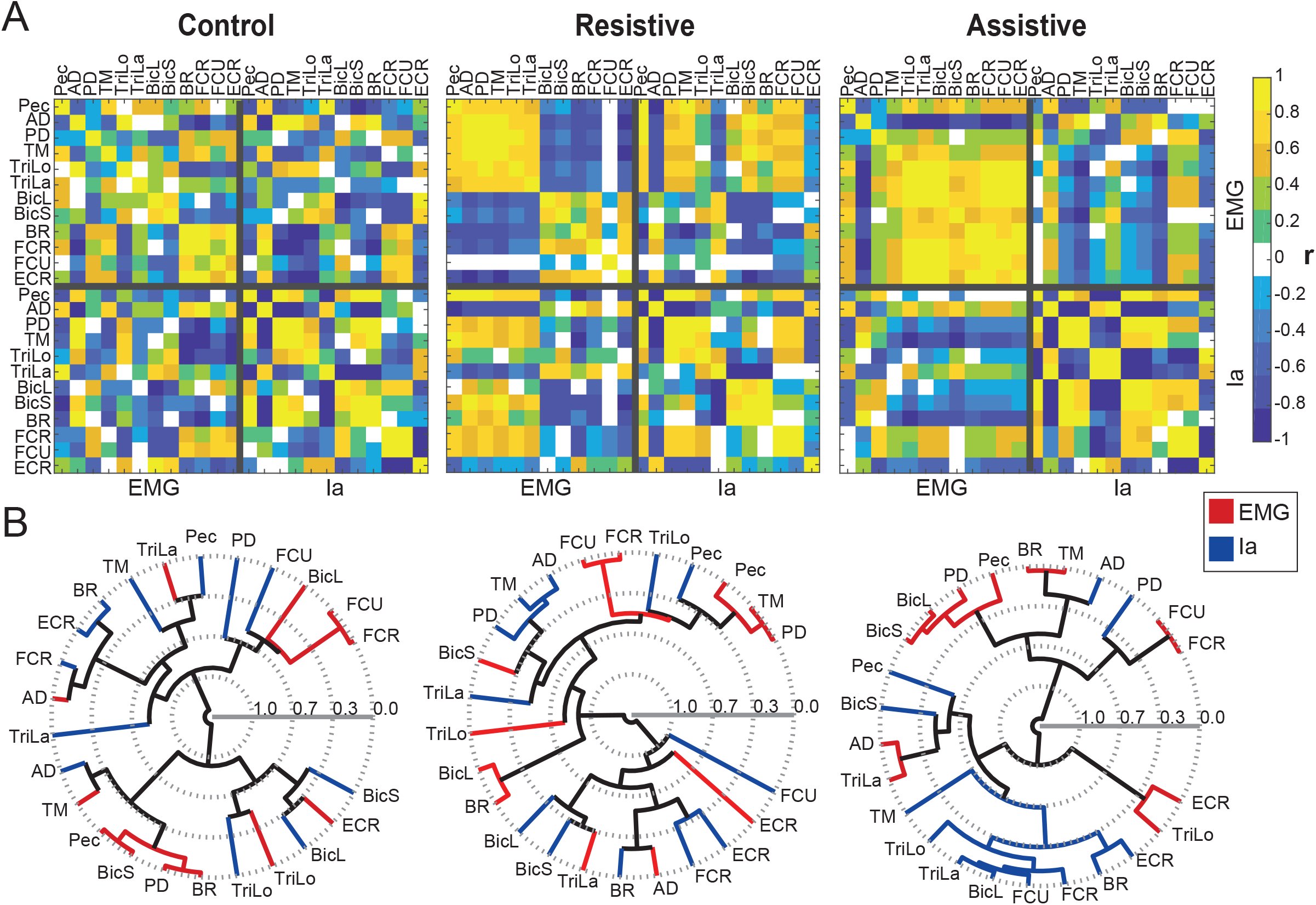
The relationships between EMG and Ia profiles per task in a representative subject. The plots are formatted as in Fig. 4. A. The correlation matrix between normalized EMG and Ia profiles for the same subject as in Fig. 4. B. Hierarchical clustering of the correlation matrix in A.

In contrast, in the Resistive task, the EMG cluster split into two, a proximal EMG cluster Pec/TM/AD/PD/TriLa/TriLo that was present in 60 % of subjects (R^2^ = 0.31 ± 0.14) and a distal EMG cluster Br/FCR/FCU/ECR that was present in 80 % of subjects (R^2^ = 0.32 ± 0.16; Fig. 4 Resistive).

These data show that the task with positive work and the least reliance on passive dynamics (Table 1, middle column) was associated with high co-contraction within two proximal and distal EMG clusters.

In the Control task, proximal muscle activity split further into smaller EMG clusters Pec/TM (R^2^ = 0.20 ± 0.23), TriLo/TriLa (R^2^ = 0.18 ± 0.15), BicS/BicL (R^2^ = 0.21 ± 0.18), while the distal EMG cluster Br/FCR/FCU/ECR (R^2^ = 0.34 ± 0.15) remained intact in 80 % of subjects (Fig. 4 Control). These data show that the largest negative work at the shoulder in the Control task (Table 1, first column) was accompanied by the lowest amount of co-contraction within proximal EMG clusters. Overall this analysis shows that the muscle groups defined by shared variance are task-dependent and that they consist of both agonistic and antagonistic muscles.

Similar to EMG, more shared variance was observed between Ia profiles in the Assistive task (R^2^ = 0.62 ± 0.14) compared to the Control and Resistive tasks (R^2^ = 0.40 ± 0.05 and R^2^ = 0.39 ± 0.09 respectively). This difference was significant (t = 4.0 and 5.4; p = 0.004 and < 0.001, Assistive/Control and Assistive/Resistive tasks respectively). This may indicate a possible contribution of primary afferents to co-contraction. However, in contrast to EMG clustering, Ia clustering was less varied between tasks. The Ia profiles clustered into smaller groups, largely based on the direction of action of the grouped muscles. These groups were Pec/PD/TM, TriLo/TriLa, BicS/BicL/Br, and FCR/FCU. The Ia groups showed high shared variance across almost all tasks (Pec/PD/TM R^2^ = 0.60 ± 0.17, 0.74 ± 0.28, and 0.67 ± 0.20; TriLo/TriLa R^2^ = 0.21 ± 0.20, 0.29 ± 0.32, and 0.72 ± 0.24; BicS/BicL/Br R^2^ = 0.65 ± 0.24, 0.75 ± 0.16, and 0.79 ± 0.11; FCR/FCU R^2^ = 0.80 ± 0.27, 0.82 ± 0.22, and 0.86 ± 0.19 for the Control, Resistive, and Assistive tasks respectively). These clusters were similar to the smallest EMG clusters in the Control task. We observed that the highest co-contraction within the distal EMG cluster in the Assistive task coincided with the highest shared variance between the agonistic subsets of the distal EMG cluster and the corresponding Ia clusters (Assistive EMG/Ia R^2^ = 0.47 ± 0.15 and 0.37 ± 0.19, Control EMG/Ia R^2^ =0.29 ± 0.14 and 0.15 ± 0.10, Resistive EMG/Ia R^2^ = 0.29 ± 0.16 and 0.17 ± 0.16, for BicS/BicL/Br and FCR/FCU respectively; Fig. 4 Assistive). Similarly, the highest cocontraction within the proximal EMG cluster in the Resistive task coincided with the highest shared variance between the agonistic subsets of the proximal EMG clusters and the corresponding Ia clusters (Resistive EMG/Ia R^2^ = 0.34 ± 0.11 and 0.24 ± 0.09, Control EMG/Ia R^2^ = 0.25 ± 0.08 and 0.16 ± 0.11, Assistive EMG/Ia R^2^ = 0.22 ± 0.09 and 0.21 ± 0.13 for Pec/PD/TM and TriLo/TriLa respectively; Fig. 4 Resistive). This shows that the small anatomical clusters observed in Ia profiles are not task dependent. They may contribute to the co-activation of agonists in a task-dependent manner.

The first null hypothesis that the similarity between Ia and EMG clusters is due to chance was not rejected. This is because the distribution of Fowlkes-Mallows indices (B) between EMG and Ia clusters was indistinguishable from the noise distribution at most cluster subdivisions (Fig. 5A, Supplementary Table 1). Some subjects showed higher-than-chance similarities between EMG and Ia clustering at 2-cluster subdivision, which represents the broad splitting of signal profiles based on gross agonist/antagonist action. However, overall this analysis shows that the small Ia clusters may be spuriously correlated with the variable EMG clusters.

**Figure 5.**
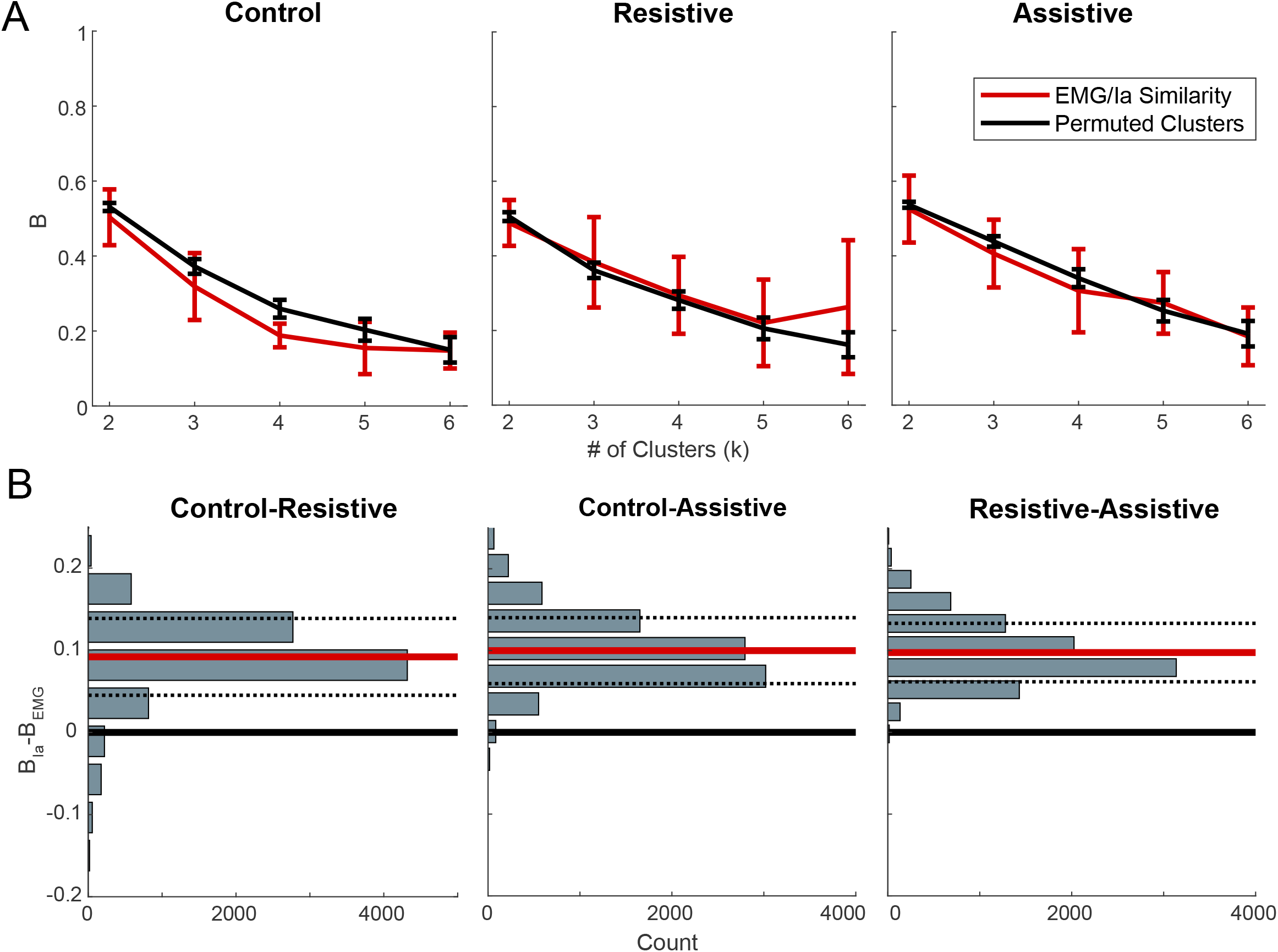
The consistency of hierarchical clustering within and across movements. A. Fowkles-Mallow Index (*B*) for the comparison between EMG and Ia cluster assignments (red) and for the comparison between EMG and permuted Ia cluster assignments representing random match (black) at different cluster subdivisions. Error bars shows pooled standard deviation across subjects. B. Histograms of the differences in *B_Ia_* and *B_EMG_* between tasks across bootstrapped hierarchal cluster trees. Horizontal thick black line is at 0, indicating no difference in *B* values between tasks; red line is at the mean of the distribution; dashed lines show standard deviations of the distributions.

The null hypothesis that Ia clusters change between tasks the same way as EMG clusters change between tasks was rejected. We have observed that the *B_Ia_* values between tasks were higher than the *B_EMG_* values between tasks (p < 0.001 for all task comparisons). This shows that between tasks the Ia clusters were more consistent than EMG clusters (Fig. 5B). This supports the observation above that the Ia clusters change less between tasks than the EMG clusters do.

In a previous computational study, we identified the functional groups of human arm muscles based on mechanical coupling using hierarchical clustering of muscle length changes across the whole range of physiological joint postures (Gritsenko et al., 2016). Here we replotted the results of hierarchical clustering of muscle lengths for the subset of muscles recorded in the current study (Fig. 6A) and compared it to the Ia clustering. We found that the muscle-length and Ia clusters were significantly more similar than expected by chance for all tasks at most cluster subdivisions (Fig. 6B, Supplementary Table 2). Thus, we have rejected the third null hypothesis that the similarity between Ia and muscle-length clusters is due to chance. This similarity was most pronounced for the Resistive movement. The similarity between muscle-length and Ia clusters further supports the results above that primary afferent feedback forms small agonistic clusters that are independent of task dynamics.

**Figure 6.**
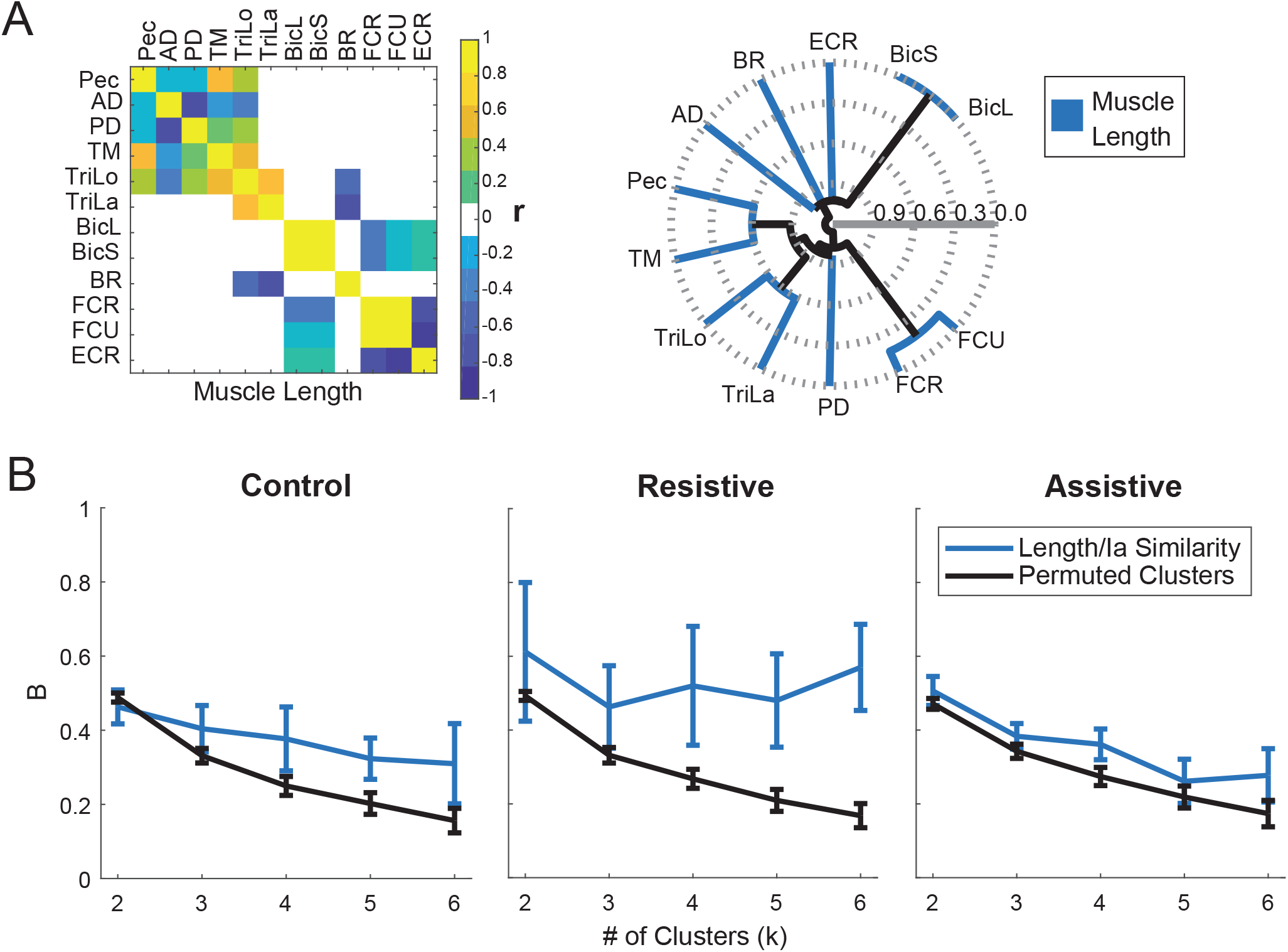
Comparison between muscle length and Ia clustering. A. The relationships between muscle length profiles per task in a default subject 0 used in (Gritsenko et al., 2016). The plots are formatted as in Fig. 4. B. The similarity between muscle length and Ia clustering for each movement. The plots are formatted as in Fig. 5A.

Lastly, we evaluated to what extent the fusimotor drive could alter Ia signal profiles to capture the features observed in muscle activity. To achieve this, we manipulated Ia model coefficients to simulate alternative fusimotor inputs, such as static set and α-γ coactivation. Models with large coefficients produced firing rates that were above those reported for human large fiber afferents (Human afferents from Malik et al. (2016): 40 imp/s; simulated afferents from Pec: 174 ± 49; AD: 331 ± 48; PD: 332 ± 49; TM: 305 ± 37; TriLo: 170 ± 25; TriLa: 243 ± 38; BicL: 171 ± 49; BicS: 177 ± 63; Br: 304 ± 93; FCR: 129 ± 42; FCU: 139 ± 40; ECR: 142 ± 35 imp/s with SD across subjects). However, the maximal simulated firing rates increased linearly with the increases in model coefficients (data not shown). Therefore, the conclusions drawn based on the data simulated at extremes using models with large coefficients will apply to the data obtained using models with lower coefficients. We found that altering the Ia model coefficients does affect the clustering pattern of Ia signals. Clusters based on the Ia model with EMG-coupled coefficients changed the most compared to those based on the original Prochazka model (Fig. 7A). However, these changes did not increase the similarity between EMG and Ia signals produced by either of the alternative Ia models (Fig. 7B and C). The last set of null hypotheses stated that the similarity between Ia and EMG clusters is not increased by alternative fusimotor drive. Neither of these hypotheses for each altered model were rejected. This shows that altering the sensitivity of muscle spindle to muscle length and its rate of change is not sufficient to reshape the primary afferent activity to reflect the dynamical features of muscle activity.

**Figure 7.**
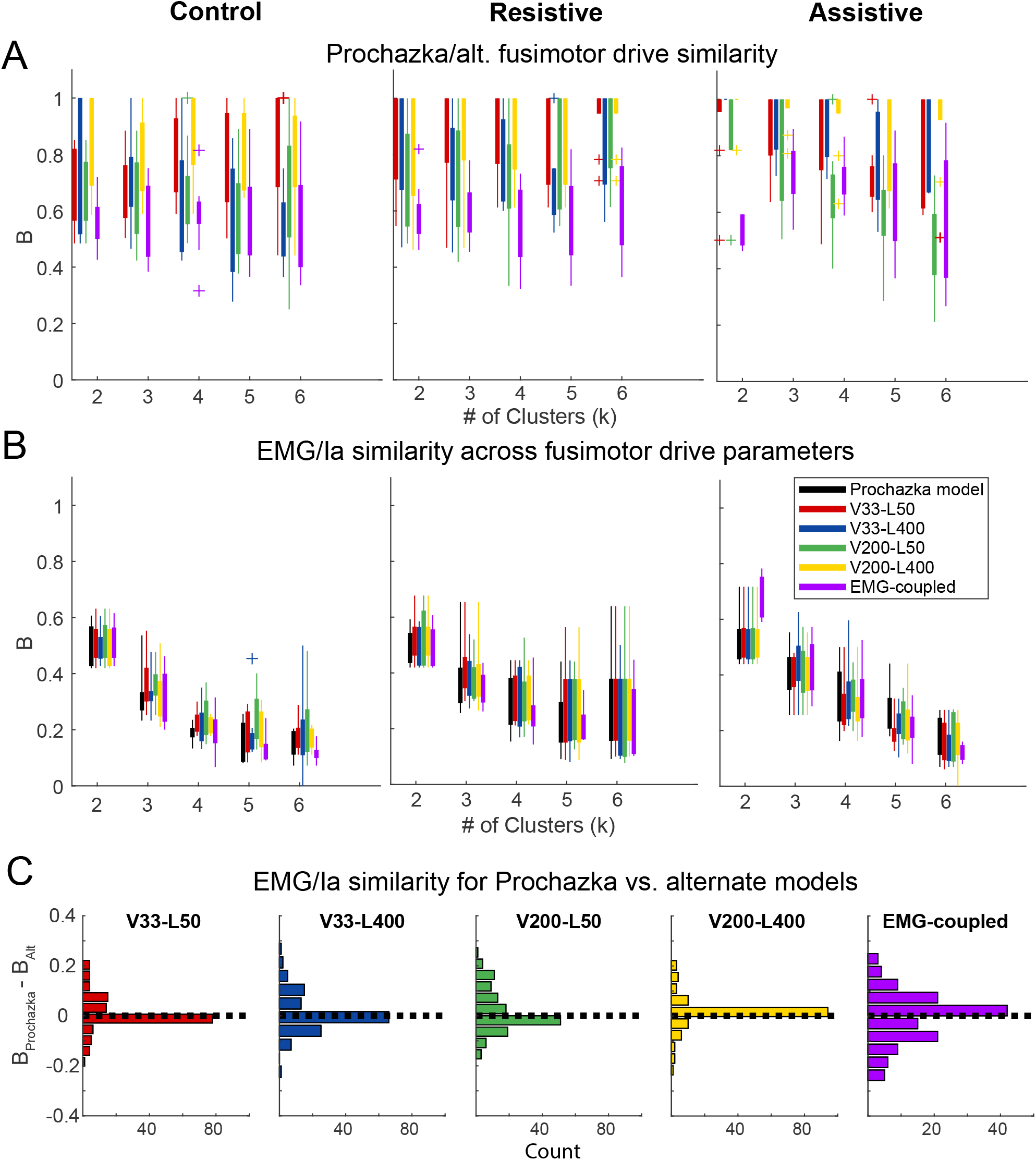
Coefficient exploration of Ia afferent firing frequency model. A. Similarity index B between models with altered coefficients and the Prochazka model across subjects. Vel. stands for velocity coefficient G1 from Eq. 2, Len. stands for muscle length coefficient G2 from Eq. 2. B. Similarity indices between EMG and Ia models with altered coefficients. Colors are the same as in A. C. Histograms of differences between indexes B of the Prochazka model and those of the alternative Ia models for individual subjects, tasks, and cluster subdivisions. Colors are the same as in A. Zero difference is close to the middle of these distributions, which indicates that no significant increases in similarity were observed across all these conditions.

## Discussion

The objectives of this study were to quantify how limb dynamics is reflected in muscle activity patterns during reaching and how primary afferent activity may help shape these patterns. We found that the VR targets defined three tasks with differential roles of passive limb dynamics that were accompanied by unique features in muscle activity. These features were in the form of flexible dynamics-dependent groups that consisted of co-activating agonistic and antagonistic muscles. We further found that the simulated primary afferent activity formed distinct groups from those formed by muscle co-activation. The Ia groups were not task dependent and formed small clusters that matched those based on musculoskeletal anatomy, i.e. flexors with flexors and extensors with extensors that span corresponding joints. Altering the sensitivity of muscle spindle to muscle length and its rate of change in simulations was not sufficient to reshape the primary afferent activity to reflect muscle activity features. Altogether, these results suggest that to solve the neuromechanical problem, the CNS controls limb dynamics through dynamics-dependent co-activation of muscles and non-linear modulation of monosynaptic primary afferent feedback (Roberts et al., 2008) (Perez et al., 2007) (Dideriksen et al., 2015).

The tasks selected for this study represent unique dynamical contexts experienced by the multisegmented limb during movement in presence of gravity. To compensate for these dynamic conditions, the forces generated by the muscles, as reflected in active muscle torques, were different in the three tasks, even when the joint motion resulting from these torques was the same (Fig. 2). Motion in the Control and Assistive tasks was produced with reliance on passive limb dynamics arising from interaction torques and gravity. The assistive action of passive dynamics in the Control task was maximally taken advantage of by the CNS, as evidenced by the lowest amount of muscle activity and co-contraction observed in EMG. However, the assistive action of passive dynamics in the Assistive task appears to have been too great. Most of the muscle action observed through muscle torques was used to decelerate the limb in the second phase of reaching (Fig. 2, Table 1). This was accompanied by the largest amount of overall co-contraction among most recorded muscles but the once spanning the shoulder (Fig. 4). This may have served to increase joint stiffness, which helped to stabilize the movement against the potentially de-stabilizing whiplash interactions between joints (Hogan et al., 1987; Burdet et al., 2001; Darainy and Ostry, 2008; Tee et al., 2010; Gritsenko et al., 2011). In contrast, motion in the Resistive task was produced against the opposing action of gravity and interaction torques between shoulder and elbow. This was accomplished with concentric contractions of two groups of proximal and distal muscles. The members of the distal group that act primarily around the wrist and hand, i.e. Br/FCR/FCU/ECR, in the Resistive task matched those from the Assistive task. The muscles that act around both shoulder and elbow did not belong to the same groups in the Resistive task compared to the Assistive task. In the Resistive task, the activity of TriLa/TriLo was more similar to the activity of muscles in the proximal group, while the activity of BicS/BicL did not consistently cluster with the activity of the rest of the muscles. Overall, our results suggest that the dynamical demands of each task define specific patterns of co-activation of agonist and antagonist muscles that form broadly-defined proximal and distal groups. These flexible task-dependent groups may reflect the neural compensation of limb dynamics (Gribble and Ostry, 1999; Gritsenko et al., 2011), providing evidence against the idea of static muscle synergies (d’Avella et al., 2003; Tresch and Jarc, 2009; Kutch and Valero-Cuevas, 2012), and supporting the idea of dynamical units of control (Sussillo et al., 2015; Yakovenko and Drew, 2015).

An interesting observation in our data is that co-contraction tended to increase with the increase in positive work at a given joint. For example, the co-contraction of the proximal EMG cluster, measured as the amount of shared variance, was high when positive work was done at the shoulder in the Resistive task compared to the other tasks. Similarly, the co-contraction of the distal EMG cluster was the highest when the most positive work was done at the elbow in the Assistive task and lower when less positive work was done in other tasks (see Results, Table 1). This provides further evidence that co-activation of agonist and antagonist muscles may constitute a neural strategy of limb dynamics compensation. Neural correlates between the activity of cells in the primary motor cortex and joint power, integral of which is joint work, have been reported in non-human primates performing similar movements (Scott et al., 2001). Also, the dynamical nature of motor cortical activity that is thought to imbed limb dynamics have been observed in non-human primates performing upper extremity movements (Churchland et al., 2012). Altogether, our results are consistent with the model of concurrent control of force and impedance demonstrated for human adaptation to tasks with altered dynamics (Franklin et al., 2008; Tee et al., 2010).

The most surprising finding of this study is that clustering of Ia profiles was distinct from that of EMG profiles and independent of the dynamical conditions of the tasks. These results suggest that primary afferent signals are providing information that is of a different modality than the outgoing activity of motoneurons. It is generally accepted that the ensemble activity of a motoneuron pool is closely linked to the force produced by the muscle innervated by it. In contrast, the primary afferent firing rate is most related to the muscle length and its rate of change (Prochazka, 1999). There is a known monosynaptic relationship between primary afferents and motoneurons innervating the same and synergistic muscles that underlies stretch reflexes, which compensate for perturbations. This anatomical arrangement with high gain, i.e. strong coupling, results in common signals that can be observed between profiles of the activity of primary afferents and the profiles of the activity of homologous motoneurons. Indeed, we observed a moderately high amount of shared variance, from 24 % to 47 %, between simulated Ia and EMG profiles, but only for small synergistic groups in tasks with high co-contraction (Fig. 5A). Furthermore, the Ia-based clusters did not change across tasks, despite the distinct kinematic and dynamic variables associated with each task. The match between Ia and EMG profiles was not improved through simulated fusimotor modulation, even with simulated α-γ coactivation (Fig. 7). This suggests that a primary afferent contribution with constant or EMG-coupled gain is not a major contributor to task-dependent co-activation of muscles observed during reaching movements. Instead, our results suggest that primary afferent feedback carries information that is orthogonal to that of muscle control signals. This indicates that the feedback signals need to be transformed into the muscle activation domain by the CNS. At the spinal level, this transformation can be accomplished through task-dependent presynaptic modulation of the monosynaptic connection between the primary afferents and *α* motoneurons (Perez et al., 2007), or through task-dependent modulation of fusimotor drive that is decoupled from motoneuron activity (reviewed in (Prochazka, 2011)). Our results with simulated fusimotor drive indicate that the neural signals that modulate primary afferent feedback need to be independent from the control signals to EMG (Fig. 7, EMG-coupled). This suggests that different type of information needs to be extracted from primary afferent feedback in order to modulate appropriately its contribution to motoneuronal activity. This information could be used for estimating limb state with the internal forward model possibly located in the cerebellum, as suggested by observations that cerebellar neuronal activity is related to arm movement kinematics (Casabona et al., 2004; Pasalar et al., 2006). The transformations required to convert sensory feedback into motor commands appropriate for the compensation for limb dynamics appear to occur at the supraspinal levels of the CNS, as evidenced by the scaling of long-latency, but not short-latency, stretch reflexes together with mechanical interactions between arm joints in humans (Kurtzer et al., 2009; Weiler et al., 2015). A control schema for sensorimotor transformation with hybrid forward/inverse models has been proposed (Wolpert and Kawato, 1998; Wolpert et al., 1998).

A potential limitation of the primary afferent model used here is its simplicity. More complex non-linear models of primary afferents exist, including models with non-linear fusimotor drive (Chen and Poppele, 1978; Mileusnic et al., 2006). These additional non-linearities may transform the Ia firing profile in such a way that it is less dependent on muscle length and its rate of change and more dependent on other signals that could be internally generated by the CNS. While our simple model cannot recreate these complex non-linearities, it does capture the basic function of the primary afferent as a length and velocity sensor. Thus, the simple model can rule out the contribution of the “raw” signal produced by such sensor to the assembly of task-dependent muscle groups.

Most muscles span at least two joints and even more DOFs, because the anatomies of most joints support motion along several rotational DOFs. For example, the shoulder joint moves along three rotational DOFs represented by functional motions of flexion/extension, abduction/adduction, and internal/external rotation. Although each DOF is independent by definition, muscle contractions would naturally couple the DOFs that belong to the joints spanned by those muscles. Thus, we would expect coupling between the shoulder and elbow DOFs because multiple muscles span both of those joints. In a prior computational study, we observed such coupling between muscle lengths across all joint excursions even without muscle contractions in a musculoskeletal model of the human arm (Gritsenko et al., 2016). We termed this mechanical coupling, as it arose purely from the anatomical arrangement of muscles on the skeleton. Interestingly, the Ia clusters observed in the current study are very similar to those identified through mechanical coupling, i.e. common signals in Ia primary afferent profiles occurred among agonist muscles that span the same joints (Fig. 6). Notably, mechanical coupling separated muscles into proximal and distal groups, where the lengths of muscles that spanned the shoulder and elbow joints did not correlate with the lengths of muscles that spanned the wrist and finger joints. This separation between proximal and distal muscles is also evident in the Ia primary afferent clusters of the present study. In contrast, the clustering of EMG signals in different movements was divided into task-dependent proximal and distal groups, which were coupled together in some tasks and decoupled in other tasks. Altogether, this suggests that one function of the sensorimotor transformation may be to couple proximal and distal muscle groups defined by anatomy using proprioceptive information about the state of each of these groups and internal predictions of the mechanical demands of the task. Future studies may test this prediction by changing the mechanical properties of proximal or distal segments and observing if the result of this perturbation will be limited to proximal or distal muscle groups or it would encompass all muscles.

## Funding

This study was supported by NIH/NIGMS U54GM104942 (RLH), T32GM081741 (RLH), T32AG052375 (RLH), Ruby Fellowship (MTB), and NIH/ NIGMS CoBRE P20GM109098 (SY, VG). The content is solely the responsibility of the authors and does not necessarily represent the official views of the NIH. This research was also sponsored by the U.S. Army Research Office and the Defense Advanced Research Projects Agency (DARPA) under Cooperative Agreement Number W911NF-15-2-0016. The views and conclusions contained in this document are those of the authors and should not be interpreted as representing the official policies, either expressed or implied, of the Army Research Office, Army Research Laboratory, or the U.S. Government. The U.S. Government is authorized to reproduce and distribute reprints for Government purposes notwithstanding any copyright notation hereon. Editorial services were supported by the West Virginia Clinical and Translational Science Institute (supported by the National Institute of General Medical Sciences, U54GM104942).

## Acknowledgements

The authors acknowledge Dr. Robert Gaunt for his valuable critical input to the interpretation of modeling results.

## Author Contributions

Substantial contributions to the conception of the work were by RH, SY, and VG; substantial contribution to the acquisition, analysis, interpretation of data, and drafting the manuscript was by RH, MB, SY, and VG.

