## Supplementary tables for "The primary afferent activity cannot capture the dynamical features of muscle activity during reaching movements"

*Supplementary Table 1: Ia and EMG  Similarity. P-values calculated as the probability that permuted cluster similarity is higher than Ia and EMG cluster similarity.*

**Control**

| Subject | Cluster 2 | Cluster 3 | Cluster 4 | Cluster 5 | Cluster 6 |
|---|---|---|---|---|---|
| 1 | 0.097 | 0.57 | 0.41 | 0.39 | 0.206 |
| 2 | 0.189 | 0.407 | 0.489 | 0.264 | 0.308 |
| 3 | 0.563 | 0.693 | 0.916 | 0.778 | 0.268 |
| 4 | 0.225 | 0.346 | 0.7 | 0.644 | 0.332 |
| 5 | 0.145 | 0.149 | 0.506 | 0.194 | 0.141 |
| 6 | 0.26 | 0.745 | 0.409 | 0.723 | 0.248 |
| 7 | 0.237 | 0.632 | 0.708 | 0.156 | 0.134 |
| 8 | 0.55 | 0.665 | 0.719 | 0.753 | 0.213 |
| 9 | 0.419 | 0.853 | 0.965 | 0.699 | 0.785 |
| Combined | 0.13927782 | 0.8353405 | 0.9668029 | 0.67177022 | 0.13464814 |

**Resistive**

| Subject | Cluster 2 | Cluster 3 | Cluster 4 | Cluster 5 | Cluster 6 |
|---|---|---|---|---|---|
| 1 | 0.493 | 0.06 | 0.113 | 0.019 | **<0.001*** |
| 2 | 0.555 | 0.508 | 0.287 | 0.586 | 0.436 |
| 3 | 0.453 | 0.153 | 0.551 | 0.444 | 0.277 |
| 4 | 0.022 | 0.003* | 0.09 | 0.153 | **0.003*** |
| 5 | **<0.001*** | 0.63 | 0.694 | 0.422 | 0.293 |
| 6 | 0.539 | 0.864 | 0.937 | 0.474 | 0.181 |
| 7 | 0.255 | 0.338 | 0.218 | 0.518 | 0.115 |
| 8 | 0.568 | 0.561 | 0.364 | 0.379 | 0.55 |
| 9 | 0.245 | 0.13 | 0.019 | 0.03 | 0.018 |
| Combined | 0.014354825 | 0.028966665 | 0.084393561 | 0.064191163 | **<0.001*** |

**Assistive**

| Subject | Cluster 2 | Cluster 3 | Cluster 4 | Cluster 5 | Cluster 6 |
|---|---|---|---|---|---|
| 1 | **<0.001*** | 0.409 | 0.469 | 0.645 | 0.898 |
| 2 | 0.148 | 0.296 | 0.729 | 0.143 | 0.134 |
| 3 | 0.533 | 0.527 | 0.572 | 0.198 | 0.215 |
| 4 | 0.218 | 0.873 | 0.859 | 0.542 | 0.089 |
| 5 | 0.521 | 0.571 | 0.912 | 0.56 | 0.322 |
| 6 | **<0.001*** | 0.092 | 0.028 | 0.15 | 0.163 |
| 7 | 0.051 | 0.286 | 0.219 | 0.457 | 0.758 |
| 8 | 0.569 | 0.848 | 0.775 | 0.37 | 0.828 |
| 9 | 0.545 | 0.258 | 0.11 | 0.007 | 0.113 |
| Combined | **<0.001*** | 0.5086354 | 0.40019888 | 0.067372203 | 0.17807019 |

*Supplementary Table 2: Ia and Muscle Length Similarity. P-values calculated as the probability that permuted cluster similarity is higher than Ia and EMG cluster similarity.*

**Control**

| Subject | Cluster 2 | Cluster 3 | Cluster 4 | Cluster 5 | Cluster 6 |
|---|---|---|---|---|---|
| 1 | 0.149 | 0.071 | 0.125 | 0.12 | 0.061 |
| 2 | 0.241 | 0.133 | 0.017 | 0.06 | 0.01 |
| 3 | 0.211 | 0.097 | 0.188 | 0.126 | 0.136 |
| 4 | 0.511 | 0.131 | 0.032 | 0.052 | 0.049 |
| 5 | 0.571 | 0.047 | 0.009 | 0.015 | 0.006 |
| 6 | 0.249 | 0.295 | 0.186 | 0.053 | 0.128 |
| 7 | 0.571 | 0.278 | 0.306 | 0.045 | 0.145 |
| 8 | 0.075 | 0.029 | 0.017 | 0.079 | **0.001*** |
| 9 | 0.072 | 0.092 | 0.015 | 0.246 | 0.126 |
| Combined | 0.087362528 | **0.0015098453*** | **<0.001*** | **<0.001*** | **<0.001*** |

**Resistive**

| Subject | Cluster 2 | Cluster 3 | Cluster 4 | Cluster 5 | Cluster 6 |
|---|---|---|---|---|---|
| 1 | 0.055 | 0.018 | 0.006 | 0.009 | **0.001*** |
| 2 | **0.001*** | **0.001*** | **0.001*** | **0.001*** | **0.001*** |
| 3 | 0.153 | 0.025 | **0.001*** | **0.001*** | **0.001*** |
| 4 | 0.232 | 0.286 | 0.059 | 0.147 | **0.001*** |
| 5 | 0.015 | 0.07 | 0.107 | 0.091 | 0.012 |
| 6 | **0.001*** | 0.007 | **0.005*** | **0.001*** | **0.001*** |
| 7 | 0.011 | 0.354 | 0.272 | **0.003*** | **0.001*** |
| 8 | 0.013 | 0.083 | 0.021 | 0.023 | 0.011 |
| 9 | 0.552 | 0.102 | 0.009 | 0.022 | **0.005*** |
| Combined | **<0.001*** | **<0.001*** | **<0.001*** | **<0.001*** | **<0.001*** |

**Assistive**

| Subject | Cluster 2 | Cluster 3 | Cluster 4 | Cluster 5 | Cluster 6 |
|---|---|---|---|---|---|
| 1 | 0.215 | 0.066 | 0.048 | 0.114 | 0.045 |
| 2 | 0.069 | 0.269 | 0.024 | 0.055 | **0.005*** |
| 3 | 0.06 | 0.261 | 0.096 | 0.449 | 0.109 |
| 4 | 0.078 | 0.269 | 0.152 | 0.156 | 0.03 |
| 5 | 0.063 | 0.088 | 0.102 | 0.447 | 0.177 |
| 6 | 0.011 | 0.037 | 0.076 | 0.126 | 0.131 |
| 7 | 0.083 | 0.273 | 0.138 | 0.167 | 0.157 |
| 8 | 0.02 | 0.145 | 0.146 | 0.431 | 0.407 |
| 9 | 0.02 | 0.161 | 0.146 | 0.118 | 0.078 |

| | | | | | |
|---|---|---|---|---|---|
| Combined | **<0.001*** | 0.009616196 | **<0.001*** | 0.030750155 | **<0.001*** |
